# Mapping sequences can bias population receptive field estimates

**DOI:** 10.1101/821918

**Authors:** Elisa Infanti, D. Samuel Schwarzkopf

## Abstract

Population receptive field (pRF) modelling is a common technique for estimating the stimulus-selectivity of populations of neurons using neuroimaging. Here, we aimed to address if pRF properties estimated with this method depend on the spatio-temporal structure and the predictability of the mapping stimulus. We mapped the polar angle preference and tuning width of voxels in visual cortex (V1-V4) of healthy, adult volunteers. We compared sequences orderly sweeping through the visual field or jumping from location to location employing stimuli of different width (45° vs 6°) and cycles of variable duration (8s vs 60s). While we did not observe any systematic influence of stimulus predictability, the temporal structure of the sequences significantly affected tuning width estimates. Ordered designs with large wedges and short cycles produced systematically smaller estimates than random sequences. Interestingly, when we used small wedges and long cycles, we obtained larger tuning width estimates for ordered than random sequences. We suggest that, ordered and random mapping protocols show different susceptibility to other design choices such as stimulus type and duration of the mapping cycle and can produce significantly different pRF results.

## Introduction

Topographic organization is a fundamental principle of the human sensory brain and the study of its properties plays a crucial role in understanding how the brain responds adaptively to properties of the environment and current goals. Important progress in brain mapping was encouraged by the introduction of population receptive field (pRF) modelling by Dumoulin & Wandell (2008). This approach aims at estimating the aggregate receptive field of all neurons within a voxel in functional magnetic resonance imaging (fMRI) scans. Essentially, a pRF identifies the location in sensory space that drives a voxel’s response, the spread of the responsive region and its shape (Dumoulin & Wandell, 2008; Silson, Reynolds, Kravitz, & Baker, 2018; Wandell & Winawer, 2015; Zeidman, Silson, Schwarzkopf, Baker, & Penny, 2018; Zuiderbaan, Harvey, & Dumoulin, 2012). A flourishing literature in the past ten years has shown that pRF modelling constitutes a powerful and sensitive approach for describing the fundamental properties of human visual cortex. The pRF size increases with increasing eccentricity and along the visual hierarchy (Amano, Wandell, & Dumoulin, 2009; Dumoulin & Wandell, 2008). Heterogeneities in pRF properties have been revealed also between different portions of the visual field (Moutsiana et al., 2016; Silson et al., 2018; Silva et al., 2018), across individuals (Moutsiana et al., 2016) and across populations (Anderson et al., 2016; Schwarzkopf, Anderson, de Haas, White, & Rees, 2014; Smittenaar, Macsweeney, Sereno, & Schwarzkopf, 2016). Moreover, pRF properties have been used to investigate neural plasticity of the visual system during development (Dekker, Schwarzkopf, de Haas, Nardini, & Sereno, 2017, 2019; Gomez, Natu, Jeska, Barnett, & Grill-Spector, 2018) or evaluate adaptive changes in the human brain resulting from diseases or trauma with pRF changes mirroring changes in visual function (Dumoulin & Knapen, 2018).

Interestingly, recent studies have also shown that pRF properties flexibly adapt to how observers engage with the stimulus. Changes in the locus of attention induce shifts in pRFs preferred location in the direction of the attended location across the entire visual field (Kay, Weiner, & Grill-Spector, 2015; Klein, Harvey, & Dumoulin, 2014; Sheremata & Silver, 2015; Vo, Sprague, & Serences, 2017). Such global changes are larger in higher visual areas (Klein et al., 2014) along both the ventral (Kay et al., 2015) and the dorsal stream (Sheremata & Silver, 2015). Moreover, recent studies indicate that pRF size and eccentricity vary in concert when the task requires to move the focus of attention from fixation to the mapping stimulus, suggesting that processing resources are adaptively redistributed to optimize the sampling of the visual stimulus according to task requirements (Kay et al., 2015; van Es, Theeuwes, & Knapen, 2018). Similarly, changes in spatial tuning of population receptive field and in their eccentricity have been observed as a consequence of changes in attentional load at fixation (de Haas, Schwarzkopf, Anderson, & Rees, 2014).

One aspect this literature has mostly overlooked is the influence of spatial predictability of visual stimuli in mapping estimates. Phase-encoded retinotopic mapping experiments (Engel et al., 1994; Sereno et al., 1995) and most pRF studies (e.g. Dumoulin & Wandell, 2008; Harvey & Dumoulin, 2011; Moutsiana et al., 2016; van Dijk, de Haas, Moutsiana, & Schwarzkopf, 2016; Yildirim, Carvalho, & Cornelissen, 2018 with few exceptions, e.g. Binda, Thomas, Boynton, & Fine, 2013; Kay et al., 2015; Thomas et al., 2015) typically employ ordered stimulus sequences - such as rotating wedges, contracting and expanding rings, or sweeping bars - to map visual areas. In such designs, the orderly presentation of the stimulus carries an inherent spatiotemporal regularity in the mapping sequence. Such regularity has two main consequences: 1) the predictability of the stimulus location, 2) the systematic consecutive stimulation of adjacent spatial locations.

Both consequences could result in fMRI responses beyond the directly stimulated voxels. Specifically, the position of a coherently moving stimulus can be anticipated based on its current location and the direction of motion. The predictability of the stimulus location could induce an anticipatory response in such locations (Ekman, Kok, & de Lange, 2017). Moreover, knowledge of the upcoming stimulus location can provide spatial cues to direct attention to the relevant portion of the screen affecting pRF estimates accordingly (Kastner, Pinsk, De Weerd, Desimone, & Ungerleider, 1999). On the other hand, the consecutive stimulation of adjacent locations in space can generate a ‘‘traveling wave’’ of activity across the cortical surface that would cause the BOLD signal to spread across neighboring voxels (Engel et al., 1994). The permeability of pRF estimates to spatiotemporal properties of the sequences has important implications also for the reliability of the estimated parameters.

In a study aiming to minimize biases when measuring visual cortex reorganization, Binda and colleagues (2013) compared pRF estimates using ordered sequences (i.e. sweeping bars) and m-sequences of multifocal stimuli. The multifocal method consists in the presentation of multiple visual stimuli presented at different locations designed to minimize the spatiotemporal correlation of visual stimulation (Vanni et al., 2005). They fitted a standard 2D-Gaussian model to voxel responses and observed that pRF size estimates (σ) in areas V1-V3 were systematically larger when ordered mapping sequence were employed. The authors suggested that differences in the mapping sequence can lead to different pRF estimates, but they did not directly address the distinctive impact of expectations and spatiotemporal regularities. Moreover, in this study the two mapping protocols differed not only in their spatiotemporal sequence dependencies, but also in stimulus shape and size, field coverage, and scanning protocol.

In this study, we aim to characterize to what extent spatiotemporal regularities in the mapping sequence affect the pRF parameter estimates in visual cortex, disentangling the role of spatial expectations and the impact of non-linear summation of the BOLD signal when adjacent locations are stimulated over a short interval. We employed functional MRI and a pRF mapping approach (Dumoulin & Wandell, 2008) to estimate the polar angle preference and the tuning response of voxels in visual cortex. We tested the same participants in three fMRI experiments using mapping sequences that differed in the spatial contingencies of consecutive wedge stimuli and in their predictability: ordered (rotating clockwise or anticlockwise), predictable, and unpredictable. In addition, we compared sequences employing stimuli of different width (wedge angle of 45deg vs 6deg) that covered the entire visual field in cycles of variable duration (9s vs 60s). We modelled pRFs of polar angle as a circular Gaussian tuning function with two parameters: the polar angle preferred response and its spread quantified as full-width half-maximum (FWHM). We compared polar angle estimates and tuning functions of pRFs in functionally defined occipital ROIs (V1, V2, V3, V3A, V4) based on the individual maps obtained from an independent mapping experiment using typical methods. Finally, we compared empirical results and simulated data as an aid for understanding the biases and reliabilities of pRF estimates.

Results suggest that the spatiotemporal regularities in the mapping protocol significantly affected pRF size (tuning width) estimates in agreement with what was previously observed for pRF size in the visual (Binda et al., 2013) and the auditory domain (Thomas et al., 2015). Moreover, we observed that the direction of the effect depended on the duration of the mapping cycle. Our results, however, do not indicate any reliable influence of stimulus predictability on pRF properties. Finally, we observed that while the ordered sequence led to the highest goodness of fit, the parameters estimated in this condition were not superior to those obtained with random conditions.

## Experiment 1

Here we asked whether the spatiotemporal structure of mapping sequences used in retinotopic mapping experiments influences the resulting parameter estimates. In particular, we tested whether pRF parameters depend on the subsequent stimulation of adjacent locations that characterize ordered mapping protocols by contrasting an *ordered rotating condition* with a random one. We further tested the hypothesis that such effects on parameter estimates depend on the predictability of the stimulus location by contrasting a predictable, non-ordered, condition with a random one.

### Materials and methods

#### Participants

Five experienced participants took part in two sessions of the experiment (1 author; age range: [24-35]; 4 females; one left-handed). All participants had normal or corrected-to-normal visual acuity. Participants gave their written informed consent to take part in the study and were financially compensated for their participation. All procedures were approved by the University College London Research Ethics Committee.

#### Stimuli and Task

Stimuli were presented using a custom MATLAB script (Mathworks Inc., Massachusetts, USA) and the Psychophysics Toolbox 3.8 (Brainard, 1997; Pelli, 1997). They were projected on a screen (1920 x 1080 pixels; 36.8 x 20.2 cm) at the back of the scanner bore and presented by means of a mirror mounted on the head coil at a total viewing distance of approximately 68 cm.

The mapping stimulus was a discretely moving wedge-shaped aperture that showed coloured natural images (1080 x 1080 pixels) depicting landscapes, textures, animals, faces, or pieces of writing randomly redrawn every 500 ms and presented on a mid-grey background. The wedge aperture extended from 0.38 to 8.5 degrees of visual angle (dva) in eccentricity. Each wedge aperture subtended 45° in terms of polar angle and was centred at one of eight polar angles (0°, 45°, 90°, 135°, 180°, 225°, 270°, 315°) dividing the circle in 8 non-overlapping locations (Figure 1A). The centre of the wedges was shifted by 15° in separate runs in order to increase the spatial granularity of the mapping.

**Figure 1.**
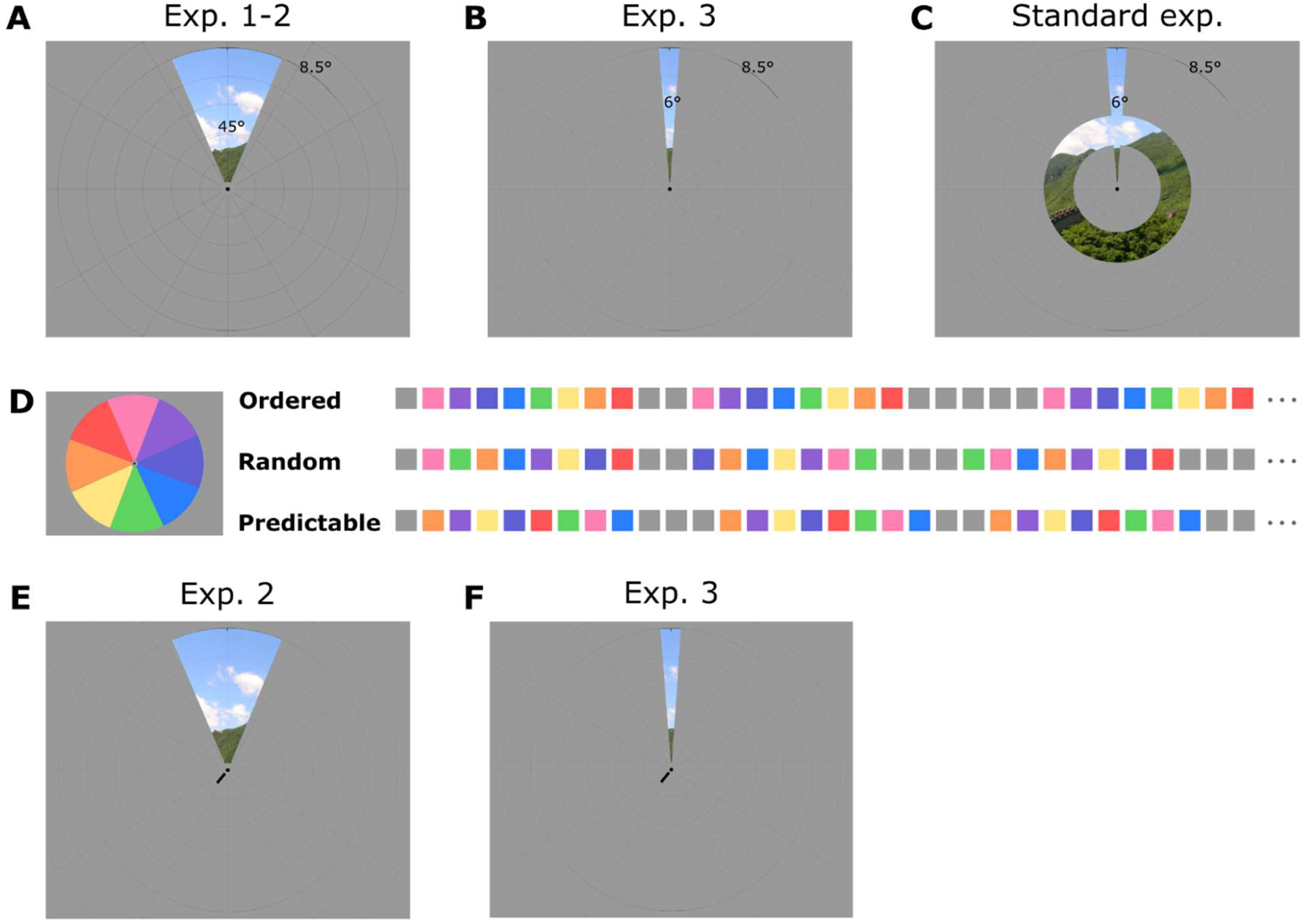
Stimuli and mapping sequences for Experiments 1-3 and the standard 2D mapping experiment. A) Large wedge mapping stimuli used in Experiments 1-2, B) thin wedge mapping stimulus used in Experiment 3, and C) wedge-and-ring mapping stimulus used in the Standard experiment. D) Spatiotemporal structure for ordered, random and predictable sequences in Experiment 1, each square represents 1 s and the colour denotes the polar angle (see colour wheel). E-F) Central cue used in predictable and random-cue sequences in Experiment 2 and 3.

Participants were instructed to continually maintain their gaze on a central fixation dot with a diameter of 0.13° while covertly monitoring the movement of the mapping stimuli in the surround. To ensure that both requirements were met, we used a dual-detection task in which participants had to report whether the colour of the fixation dot turned red (fixation task) and whether an Anderson tartan pattern was presented on the wedges (image detection task) (Moutsiana et al., 2016; van Dijk et al., 2016). To aid participant’s compliance with fixation requirements, a low contrast polar grid (line width of 0.02°; opacity of 10.2%) composed of 10 circles (radii evenly spaced between 0.13° and 15.14°) and 12 evenly spaced radial lines extended from fixation to the edges of the screen was superimposed onto the grey foreground and stimuli at all time. Eye movements were further monitored by means of an MRI-compatible SR Research EyeLink 1000 eye tracker.

#### Mapping sequences

Three mapping conditions were presented to each participant: *ordered, predictable* and *random* (i.e. unpredictable) (Figure 1D). In the ordered runs, the wedge rotated around fixation either clockwise or anticlockwise starting randomly at one of the 8 locations. The same direction of motion and the same starting location was maintained within each run. In the predictable runs, the wedge was presented at the 8 locations according to a predefined pseudorandomized order such that no adjacent locations were stimulated one after the other. The sequence started at a random location in different runs, but the same starting point and the same order were maintained throughout the run. Six maximally distinctive sequences were selected for each participant, three for each session. In the unpredictable runs, wedges were presented at the 8 locations in pseudorandom order (no adjacent locations could be presented in a row) and from a random starting point. A different, randomly generated, sequence was presented in each cycle (Figure 1D). For all conditions, each step of the wedge was presented for 1 s such that an entire cycle was completed in 8 s. The wedge completed 16 cycles in each run. Cycles were separated by fixation intervals of variable duration pseudo-randomized to range from 1 to 8 s in discrete steps of 1 s. Before entering in the scanner, participants performed a 30-minute task to familiarize themselves with the predictable sequences that they would encounter during the scanning session. Each sequence was presented in a separate block. Each block started with a presentation of the 8-steps sequence, presented for 7 times, after which we introduced a violation of the location order in the sequence. The participant’s task was to detect this violation of regularity and report it with a button press. Each sequence was presented in 6 consecutive blocks and was presented 20 times per block (9 correct sequences and 11 sequences with violations). In the scanner, a familiarization block preceded each mapping run in order to familiarize participants with the sequence that they would encounter during the following scanning run. Similar familiarization blocks were repeated before each run of the ordered and random conditions. For both the ordered and the predictable condition, participants performed a sequence violation-detection task in which they reported when a wedge appeared in an unexpected location according to the learned sequence (predictable condition) or the direction of motion (ordered condition). In the random condition, participants performed a 2-back task in which they reported when a stimulus was presented in a location that was occupied 2 stimuli before.

Reference retinotopic maps were obtained for each participant in an additional experiment using a combined wedge-and-ring aperture (Figure 1C) similar to what has been reported in previous studies (Alvarez, de Haas, Clark, Rees, & Schwarzkopf, 2015; Moutsiana et al., 2016; van Dijk et al., 2016). The wedge aperture extended up to 8.5 degrees of visual angle in eccentricity and subtended 12° rotating either clockwise or counter-clockwise in 60 discrete steps (1 step/s, 6° overlap between consecutive wedges). The ring aperture expanded or contracted in 36 logarithmic steps while keeping the radius of the inner circle 56-58% of that of the outer ring (minimal radius of 0.48 dva, 1 step/s, ∼90% overlap between consecutive rings). The mapping stimulus showed coloured natural images or phase-scrambled versions of them that changed every 500 ms and appeared on a mid-grey background. The type of image (intact vs phase-scrambled) alternated every 15 s. The images and the wedge-and-ring aperture were centred on a central black fixation dot (diameter: 0.13 degrees in visual angle) which was superimposed onto central disk (diameter: 0.38 degrees in visual angle). Also, a low contrast polar grid was superimposed on the stimulus. As for the previous experiments, participants performed a dual-detection task (fixation task and image detection task) while maintaining fixation on the central fixation dot.

The mapping experiment consisted of 3 runs. The wedge-and-ring aperture was presented in four blocks of 90 s (1.5 cycles of wedge rotation; 2.5 cycles of ring expansion/contraction) interleaved with a 30 s blank interval. The order of aperture movement in each run was first clockwise and expanding, then clockwise and contracting, anticlockwise and expanding, or anticlockwise and contracting.

#### Data acquisition

We acquired functional and anatomical scans using a Siemens Avanto 1.5 T MRI scanner with a customized 32-channel head coil located at the Birkbeck-UCL Centre for Neuroimaging. The two anterior channels were removed from the front half of the coil to allow unrestricted field of view leaving 30 effective channels.

Functional images were acquired using a T2*-weighted 2D echo-planar images multi-band (Breuer et al., 2005) sequence (TR = 1 ms, TE = 55 ms, 36 slices, flip angle = 75°, acceleration = 4, FOV = 96 × 96 voxels) at a resolution of 2.3 mm isotropic voxels. Each functional scan consisted of 222 acquisitions. Data were collected in two sessions (performed on consecutive days or one day apart) of 9 runs each taking approximately 90 minutes^1^. Each condition was repeated in 3 separate runs in each session. The order of runs was pseudo-randomized, with all conditions repeated every 3 runs. The ring-and-wedge mapping procedure was acquired in a separate session using the same protocol, for a total of 3 runs and 490 volumes per run.

A T1-weighted anatomical magnetization-prepared rapid acquisition with gradient echo (MPRAGE) image was acquired in a separate session (TR = 2730 ms, TE = 3.57 ms, 176 sagittal slices, FOV = 256 × 256 voxels) at a resolution of 1 mm isotropic voxels.

#### fMRI pre-processing

The data were pre-processed using SPM12 (www.fil.ion.ucl.ac.uk/spm, Wellcome Centre for Human Neuroimaging, London, UK). The first 10 volumes of each run were discarded to allow the signal to reach equilibrium. Functional images were intensity bias-corrected, realigned to the mean image of each run and then co-registered to the structural scan. All further analyses were performed using custom MATLAB code. The time series for each voxel in each run were linearly de-trended and z-score normalized. Finally, all runs belonging to the same condition were concatenated before further analyses whereas in the wedge-and-ring standard mapping experiment, the time series were averaged before the fitting analysis to increase signal to noise ratio. Functional data of each participant were projected on a surface reconstruction of the grey white matter surface estimated with FreeSurfer (Dale, Fischl, & Sereno, 1999; Fischl, Sereno, & Dale, 1999) by finding the voxel at the medial position between the grey-white matter boundary and the pial surface for each vertex in the mesh (using custom made Matlab scripts and SamSrf toolbox). All the following analyses were performed at the surface level. The same procedures were adopted for Experiment 2 and 3.

#### pRF estimates

The data from the different protocols were used to obtain independent estimates of the population receptive fields (pRFs) using a custom MATLAB toolbox for pRF analysis (SamSrf v5.84, https://doi.org/10.6084/m9.figshare.1344765.v24). For all mapping sequences, we combined a binary aperture describing the position of the mapping stimuli within each scanning volume with a model of the underlying neuronal population and convolved this with a canonical haemodynamic response function (HRF) to predict the BOLD signal in each experimental condition. For the standard wedge and ring mapping sequence, the binary aperture was a two-dimensional mask (100×100) corresponding to the stimulus location on the screen at each time point. For the main experiment, the binary aperture was a vector mask (1×360) indicating the polar angle coordinates corresponding to the mapping stimulus at each time point.

For the standard wedge and ring mapping sequence, we estimated the position and size of pRFs using a two-dimensional Gaussian function (Dumoulin & Wandell, 2008). For the polar mapping experiments, we modelled pRFs using a von Mises distribution, with µ indicating the preferred polar angle of the voxel and k corresponding to the concentration of the response (for clarity k values were transformed into full width half maximum (FWHM) as an indicator of the spread of the response of each voxel). We used a coarse-to-fine approach to determine the pRF parameters to obtain the best possible fit of the predicted time series with the observed data (Alvarez et al., 2015; Dumoulin & Wandell, 2008; Moutsiana et al., 2016; van Dijk et al., 2016). The final fine fitting also included a β parameter for the response amplitude.

#### Analyses

We only analysed the fitted parameters of those vertices for which we obtained realistic estimates (k>0) and that had a goodness of fit, R^2^, higher than a critical value based on a fixed p-value (p = 10^−8^). This corresponds to R^2^>0.026 ^2^ in Experiment 1, and R^2^>0.067 in the wedge-and-ring experiment depending on the different degrees of freedom in the three experiments^3^.

The pRF estimated coordinates from the standard wedge and ring mapping experiment were used to compute polar angle and eccentricity. Using the Delineation toolbox in SamSrf, we manually delineated the regions of interest using mirror reversals in the polar angle map, and guided by the eccentricity and field-sign map (Sereno, McDonald, & Allman, 1994). The region of interests included in our analyses were V1, V2, V3, V3A and V4. We performed all the following analyses separately for each visual ROI in each individual participant. Given the small number of participants, we did not report group statistics in the main text, but we summarized the results for single subject statistics instead.

We compared the quality of the fits across the different spatiotemporal sequences. We compared the number of responsive vertices and the median goodness of fit for each individual by means of repeated paired t-tests and Wilcoxon tests. We also used a correlation analyses to evaluate the correspondence between the observed time series for each condition and the predicted response given the stimulus location and the parameter estimates for each vertex obtained with each of the mapping protocols, convolved with an HRF. To further explore the coherence of the maps obtained with different mapping conditions, we computed the vertex-wise circular correlation of polar angle estimates and the Pearson correlation of FWHM and beta estimates between conditions, separately for each visual ROI and participant. Because vertices within a ROI are not statistically independent, we calculated the inter-correlation between the time series of all ROI vertices and used this information to correct the degrees of freedom of the correlation. Specifically, we calculated all unique pair-wise correlations between vertices (note that we treated pairs of vertices that were negatively correlated as independent, i.e. r = 0). We then calculated a weight for each vertex by subtracting these correlations from 1 and averaging the values for all pair-wise comparisons of a given vertex. Thus, in theory, if the time series of all vertices were completely independent from one another, each vertex would be weighted as 1. Conversely, if all vertices were identical, they would all be weighted as 0. The sum across these weights plus 1 is therefore a weighted estimate of the sample size which we used to determine the degrees of freedom.

Moreover, we correlated the observed time courses for each condition with the predicted time course given the estimated pRF parameters for each vertex in each experimental condition. Finally, we compared the mean FWHM across conditions and ROIs using paired t-tests at the subject level (with degrees of freedom corrected for inter-correlation between time series as described above). In these analyses, we averaged FWHM across vertices encompassing different eccentricities, as our mapping stimulus did not allow differentiating responses at different eccentricities (i.e. each wedge had a fixed radius that covered the entire visual field mapped).

### Results

We obtained reliable polar angle maps with all mapping conditions for our ROIs (Figure 2). Ordered sequences provided better fits and a larger proportion of responsive vertices than the other mapping sequences in all ROIs (Mean of median R^2^ across ROIs: *M*_*ord*_ = .10, *M*_*pred*_ = .08, *M*_*rnd*_ = .08; Mean of responsive vertices across ROIs: *M*_*ord*_ = 35%, *M*_*pred*_ = 25%, *M*_*rnd*_ = 28%). Moreover, we observed better fits for higher visual areas (Figure S 1A, D). The relatively low goodness of fit obtained with this paradigm is not surprising given the high number of degrees of freedom in our paradigm (due to concatenated time series rather than averaging experimental runs) and the simplified model used for fitting. Importantly, all vertices considered for comparisons provided a fitting that met our fixed p-value criterion of 10^−8^ in all the mapping conditions. Interestingly, parameter estimates obtained with different mapping sequences performed similarly well in predicting the observed time series for all mapping protocols, with generally more robust predictions of the ordered sequences regardless of the mapping protocol used for the estimates (Figure S 1A). Consistently, correlation analyses of model fitting results (polar angle, FWHM, beta, and R^2^) revealed substantial consistency across different mapping conditions in all experiments and all visual areas tested. We observed high significant vertex-wise correlation between R^2^ (M_ord-rand_ = .79, M_ord-pred_ = .77, M_rand-pred_ = .82; Figure 3A) in different conditions for all participants and ROIs (p<.05, Bonferroni corrected for multiple comparisons). The polar angle estimates were highly robust across conditions but showed a decrease in coherence moving up in the visual hierarchy for the predictable condition (M_ord-rand_= .86, M_ord-pred_= .56, M_rand-pred_= .68; Figure 3B).

**Figure 2.**
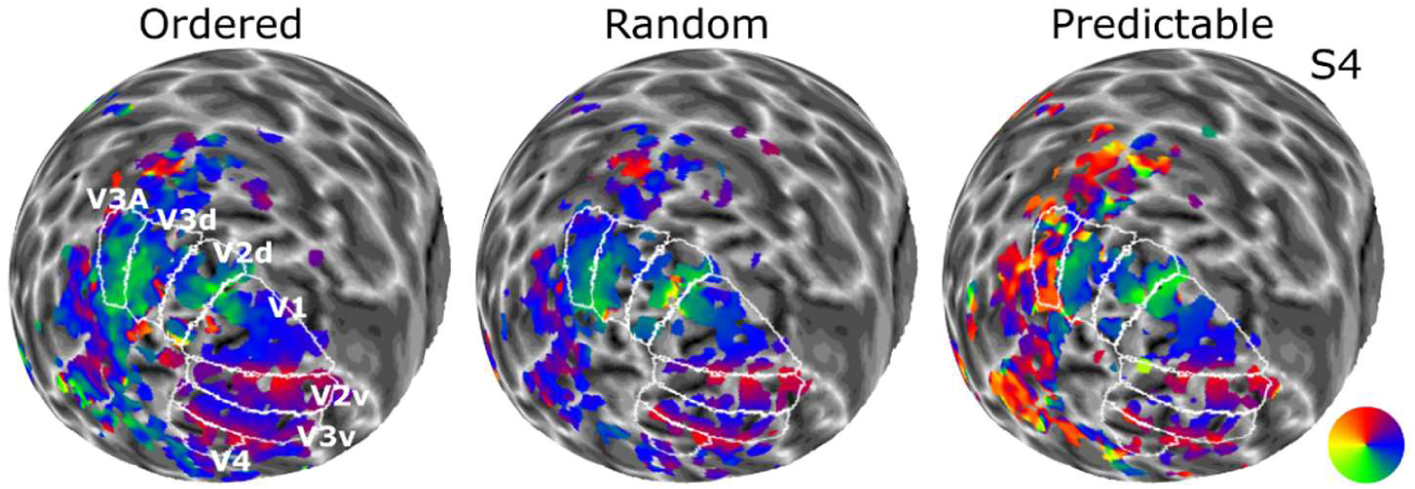
Smoothed polar angle maps for all conditions - ordered, random and predictable - in Experiment 1. In this and the following figures, images display an inflated spherical model of the left hemisphere of participant 4 (S4).

**Figure 3.**
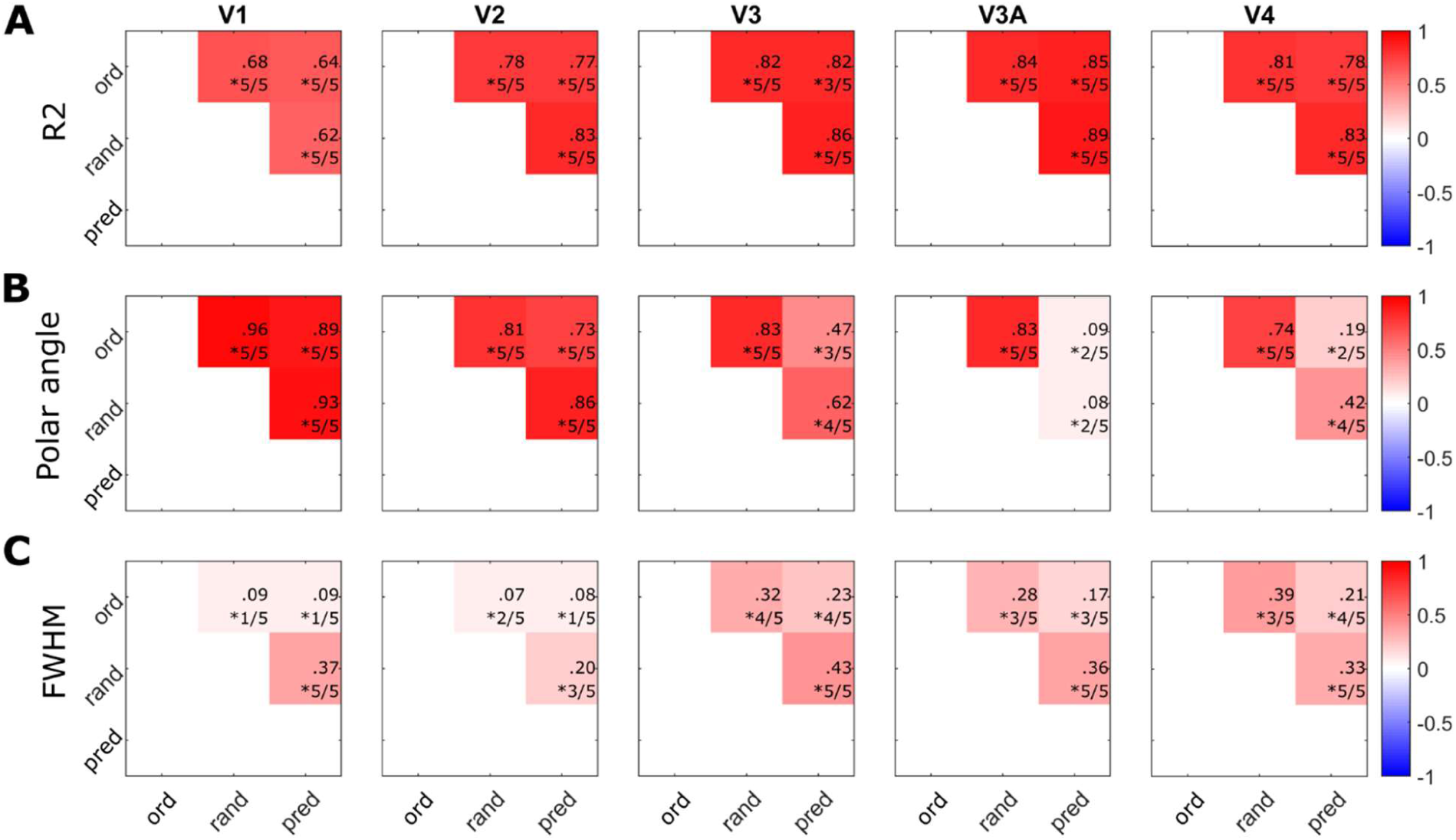
Group mean correlation matrices for A) the model goodness of fit (R^2^), B) polar angle estimates), and C) FWHM estimates in Experiment 1. Inset numbers (in this and the following figures) in each cell of the correlation matrix indicate the value of the average correlation and the proportion of participants that showed a significant correlation for each pair of conditions.

We observed positive but substantially weaker correlations for FWHM estimated, particularly in lower visual areas, V1-V2 (M_ord-rand_ = .24, M_ord-pred_ = .17, M_rand-pred_ = .35; Figure 3C).

We further explored potential biases and differences across mapping conditions and ROIs. As expected, FWHM estimates increased in the visual hierarchy. Interestingly, FWHM was also systematically influenced by the mapping sequence and a general pattern emerges for all visual areas (with the exception of V1 that shows noisier results) with results highly consistent across participants (Figure 4; significant results are reported for p<.05, Bonferroni corrected for multiple comparisons). The ordered sequence lead to significantly smaller FWHM estimates than the random sequence for most of the participants and ROIs (Figure 4B). We found similar differences between ordered and predictable sequences, although one participant showed a significant difference in the opposite direction. Interestingly, we measured smaller FWHM for predictable than random sequences (results are clearer for V2, V3 and V4).

**Figure 4.**
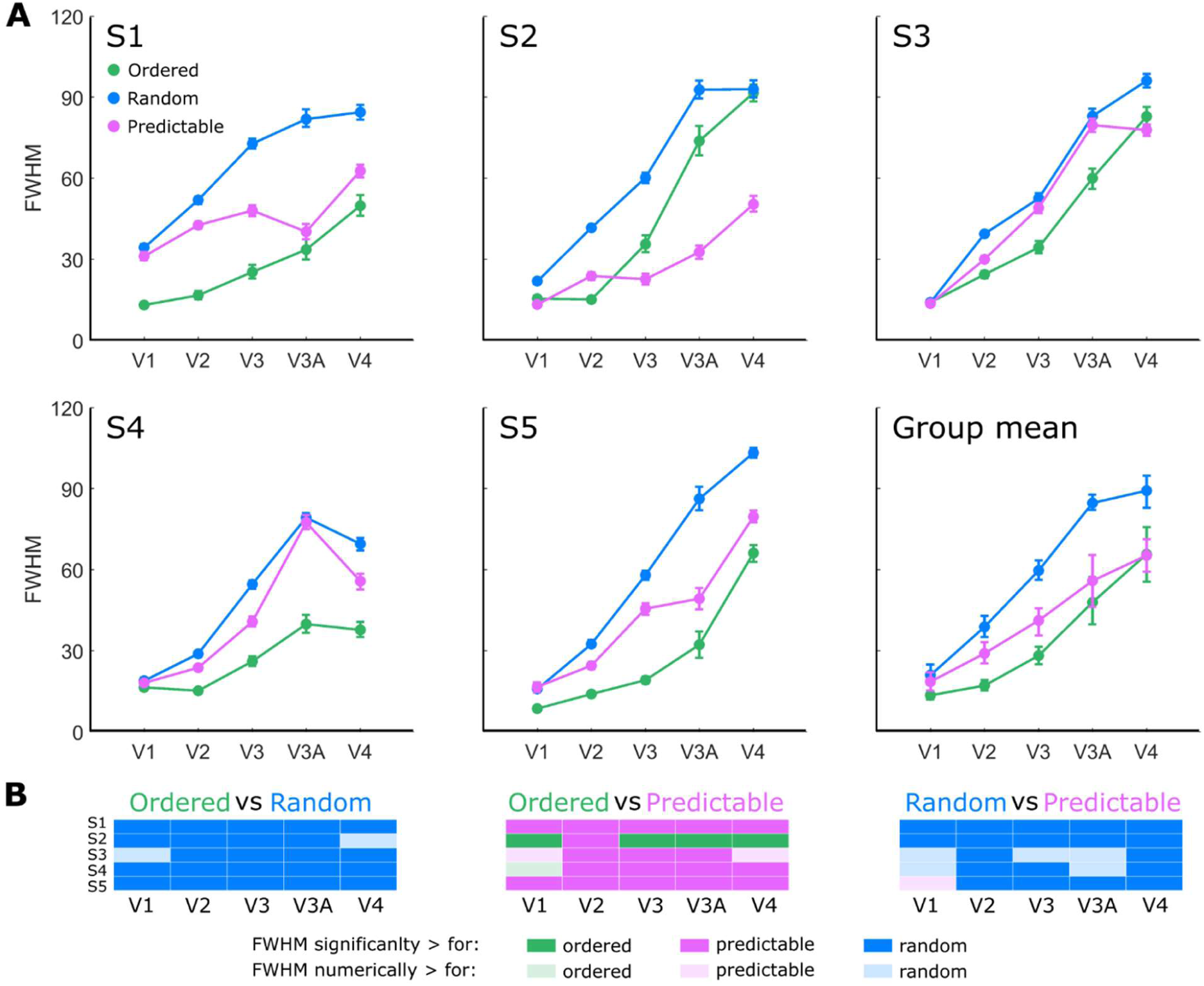
Polar angle tuning width estimates in Experiment 1 (measured as Full-Width Half-Maximum). A) Individual and group mean FWHM estimates for different mapping sequences and visual areas. In this and the following figures, error bars in both the individual and group plots represent bootstrapped 95% confidence intervals. B) Visualisation of single subject statistics (p<.05 corrected for multiple comparisons).

## Experiment 2

Experiment 1 revealed systematic differences in polar angle tuning functions estimated with different mapping protocols. This suggests that predictability might influence the tuning response of population of neurons in visual cortex. However, the predictability of our mapping sequences depended on the repetition of fixed spatiotemporal structure throughout each run. Such idiosyncrasies in the predictable sequences could have been responsible for relatively narrow tuning width estimates (Figure 4) and the poorer agreement in polar angle estimates of this sequence with the ordered and the random one (Figure 3B).

To address whether the fixed spatiotemporal structure of the predictable sequence was responsible for the observed results, we repeated Experiment 1, but this time creating a predictable sequence that was structurally indistinguishable from the random one. Rather than using a repeated sequence, we rendered the sequence predictable by the use of a small visual cue. We then compared the tuning width response of this sequence with the random, non-predictable one.

### Materials and methods

#### Participants

Four of the original subjects took part in the two sessions of Experiment 2 (one author; age range: [24-35]; 3 females). All participants had normal or corrected-to-normal visual acuity and gave their written informed consent to participate to the experiment as in Experiment 1.

#### Stimuli and Mapping sequences

Experiment 2 was set up with the same apparatus and mapping stimuli used in Experiment 1 (Figure 1A). We compared four mapping sequences - ordered, predictable and two random ones. Crucially, we changed how we induced the predictability of the wedge location in the predictable condition. The predictable and random sequences were generated using the same algorithm, i.e. wedges were presented at different locations in pseudorandom order with no adjacent locations presented in a row. In contrast to Experiment 1, we generated a different sequence in each cycle thus completely matching the spatiotemporal structure of random and predictable sequences. We maintained the difference in predictability of the wedge locations by means of a centrally presented oriented line that cued the location of the wedges (Figure 1E). The cue (0.33 x 0.07 degrees in visual angle) extended from the centre of the screen. It appeared 200 ms before the onset of each wedge stimulus and remained on the screen for 200 ms (Figure 1C). In the *ordered* and the *predictable* conditions, the cue pointed towards the centre of the upcoming wedge. The two random conditions were both unpredictable but differed for the presence or absence of the central cue. In the random condition with non-predictive cue (*random-cue*), the cue pointed to the location of the previous wedge. In the random condition without cue (*random-no cue*), no cue was presented. Thus, neither of the random conditions contained any information about the location of the upcoming wedge.

For all conditions, each step of the wedge was presented for 1 s such that an entire cycle was completed in 8 s. Cycles were separated by fixation intervals of variable duration ranging from 1 to 8 s in steps of 1 s. Each functional scan consisted of 303 acquisitions. Data were collected in two sessions (performed on consecutive days or one day apart) of 12 runs each taking approximately 90 minutes. Each condition was repeated in 3 separate runs in each session. All conditions were presented in randomized order every 4 runs.

#### Analyses

As in Experiment 1, we only analysed the fitted parameters of those vertices for which we obtained realistic estimates (k>0) and that had a goodness of fit, R^2^, higher than a critical value based on a fixed p-value (p = 10^−8^). This corresponds to R^2^>0.019 in Experiment 2.

### Results

Experiment 2 mostly replicated the results of Experiment 1 with even clearer polar angle maps (Figure 5) and higher consistencies between parameter estimates (Figure 6), possibly due to the higher number of volumes collected.

**Figure 5.**
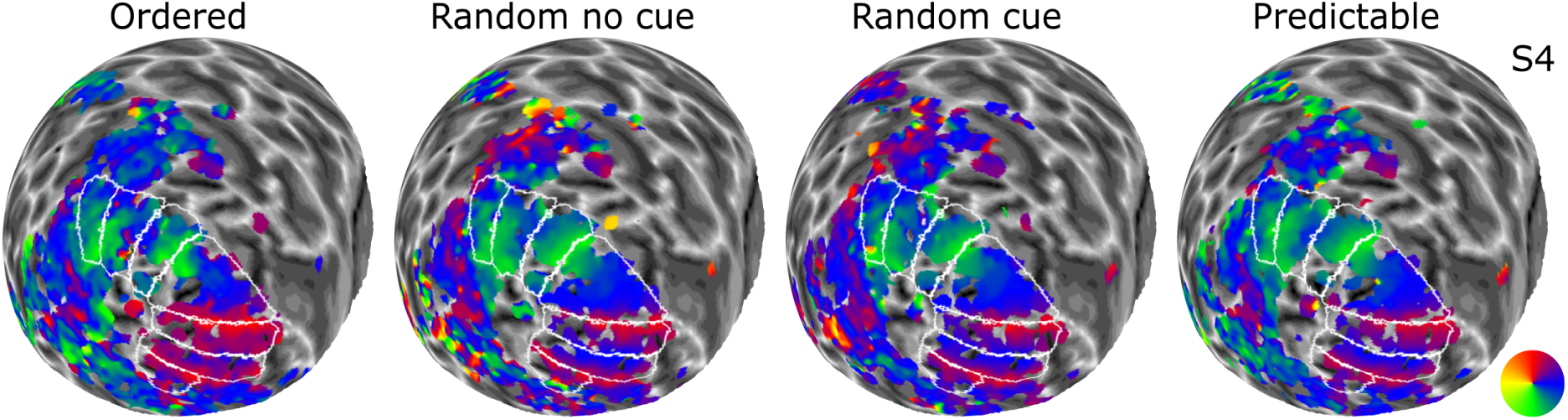
Smoothed polar angle maps for all conditions - ordered, random with or without uninformative cue, and predictable - in Experiment 2.

**Figure 6.**
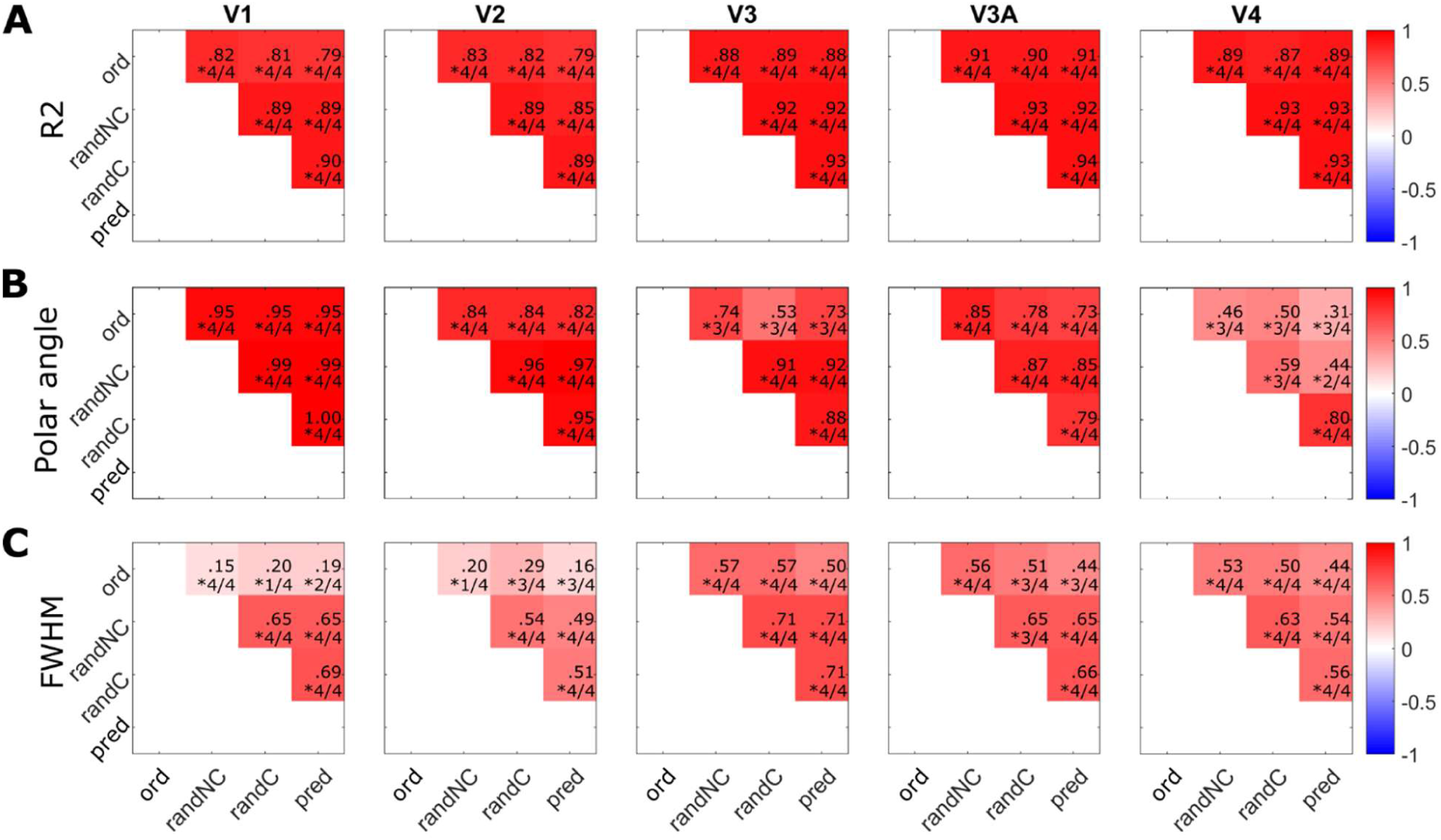
Mean correlation matrices for A) goodness of fit (R^2^). B) polar angle, and C) FWHM estimates in Experiment 2.

Ordered sequences provided better fits to the data than predictable and random sequences (Figure S 1B, E; Mean of median R^2^ across ROIs: *M*_*ord*_ = .09, *M*_*pred*_ = .08, *M*_*randNC*_ = .08, *M*_*randC*_ = .08; Mean of responsive vertices across ROIs: *M*_*ord*_ = 43%, *M*_*pred*_ = 41%, *M*_*randC*_ = 40%, *M*_*randNC*_ = 37%). However, our analyses confirmed that the parameters estimated in the different conditions performed equally well in predicting the time series data (Figure S 2B).

We also replicated differences in FWHM estimates with different sequence structures with ordered sequences leading to systematically smaller FWHM estimates than random and predictable sequences (p<.05 corrected, for all participants and ROIs but one comparison for S2 V3A as illustrated in Figure 7**Error! Reference source not found**.). Importantly, we did not observe any systematic differences between predictable and random sequences with the exception of V2, where FWHM were systematically smaller for predictable than random sequences as also shown in Experiment 1 (significant difference for all participants in the comparison with the random-no cue condition and with all participants but one in the random-cue condition).

**Figure 7.**
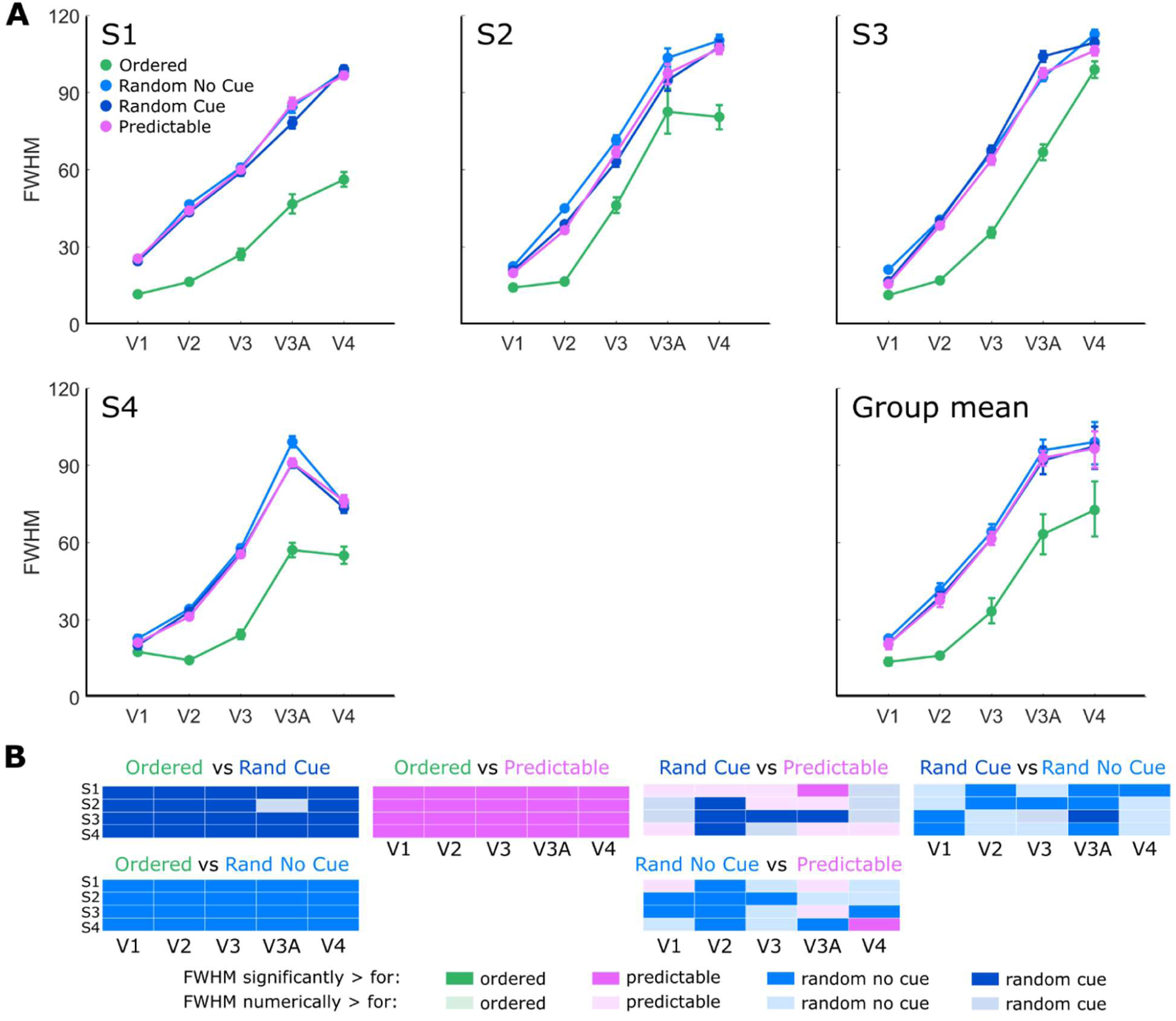
Polar angle tuning width estimates in Experiment 2. A) Individual and group mean FWHM estimates. B) Visualisation of single subject statistics (p<.05 corrected for multiple comparisons).

## Experiment 3

Experiment 2 suggested that the spatiotemporal structure of the mapping sequence, rather than its predictability is responsible for the differences in FWHM estimates. The finding that ordered sequences yielded narrower tuning widths than random sequences in both Experiment 1 and 2 contrasts with previous studies. A comparison of orderly moving bars and multifocal stimuli revealed the opposite pattern of results, with the largest pRF estimates obtained with ordered sequences (Binda et al., 2013). However, it is not clear whether pRF size estimates might have been affected by surround suppression of response in the multifocal stimuli (Pihlaja, Henriksson, James, & Vanni, 2008). Moreover, our results might be affected by the short mapping sequences we adopted. To address this hypothesis, we replicated Experiment 2 with the same mapping conditions and the same participants, but we varied the size of the mapping stimulus as well as the duration of the mapping cycle.

### Materials and methods

#### Participants

The same four participants (including one author) that took part in Experiment 1 and 2 participated also in both sessions of Experiment 3. All participants gave their written informed consent to participate to the experiment.

#### Stimuli and Mapping sequences

The mapping stimulus in Experiment 3 was a discretely moving wedge aperture subtending 6° and dividing the circle in 60 non-overlapping locations, no shifts were introduced across runs (Figure 1B). The mapping sequences used in Experiment 3 were generated in the same way as those in Experiment 2 resulting in four experimental conditions: *ordered, predictable, random-no cue*, and *random-no cue* (Figure 1F). The distinctive difference between Experiment 2 and 3 is only the aperture size and, consequently, the duration of the mapping cycle. For all conditions, each step of the wedge was presented for 1 s such that an entire cycle was completed in 60 seconds (60 wedges of 6°, 4 cycles). Cycles were separated by fixation intervals of variable duration ranging from 1 to 8 s in steps of 1 s. Each functional scan consisted of 295 acquisitions whilst other scanning details remained identical to Experiment 2.

#### Analyses

As in the previous experiments, we only analysed the fitted parameters of those vertices with k>0 and goodness of fit, R^2^>0.019 (based on fixed p-value p = 10^−8^).

### Results

Parameter estimates were consistent across mapping sequences (Figure 8, Figure 9; Figure S 2C).

**Figure 8.**
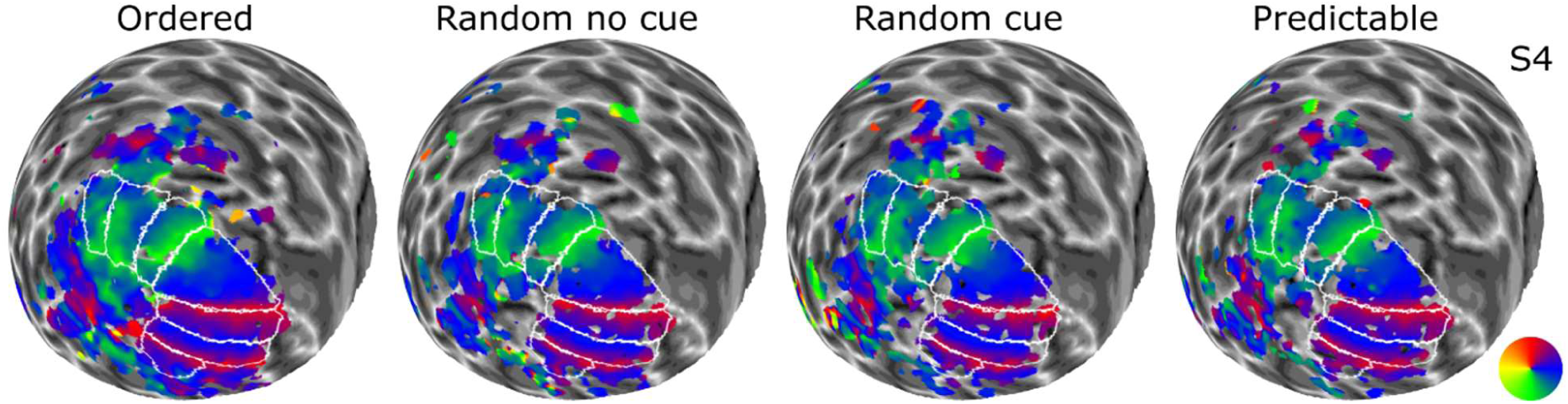
Smoothed polar angle maps for all conditions - ordered, random with or without uninformative cue, and predictable - in Experiment 3.

**Figure 9.**
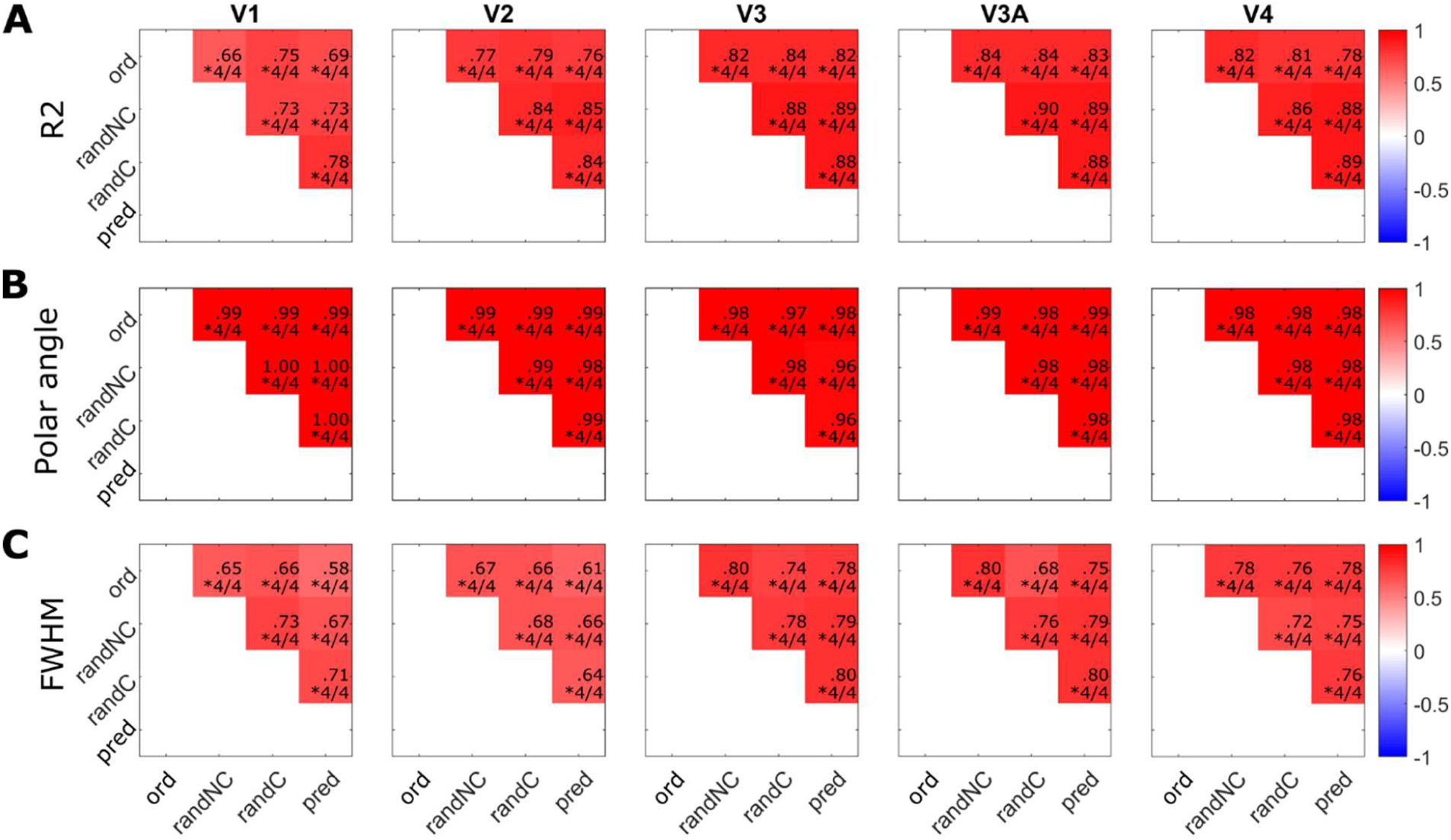
Mean correlation matrices for A) goodness of fit (R2). B) polar angle, and C) FWHM estimates in Experiment 3.

Ordered sequences provided much better fits to the data than predictable and random sequences (Mean of of median R^2^: *M*_*ord*_ = .14, *M*_*pred*_ = .06, *M*_*randNC*_ = .06, *M*_*randC*_ = .06; Mean of responsive vertices across ROIs: *M*_*ord*_ = 61%, *M*_*pred*_ = 37%, *M*_*randC*_ = 38%, *M*_*randNC*_ = 38%; Figure S 1C, F). However, the parameters estimated in the different conditions performed equally well in predicting the time series data and they all yielded the best results when predicting the ordered sequence (Figure S 2C).

Experiment 3 produced an interesting inversion of the pattern of results in terms of FWHM estimates (Figure 10) compared to the previous experiments. Although there was no consistent pattern in the contrast between predictable and random sequences, FWHM estimates were systematically larger for ordered than random or predictable sequences for V3, V3A and V4. Similar results were found for V2 but with less consistent results across participants. Results for V1 were less clear but seem to suggest the opposite: FWHM were smaller for estimates obtained with an ordered rather than a random or predictable sequence.

**Figure 10.**
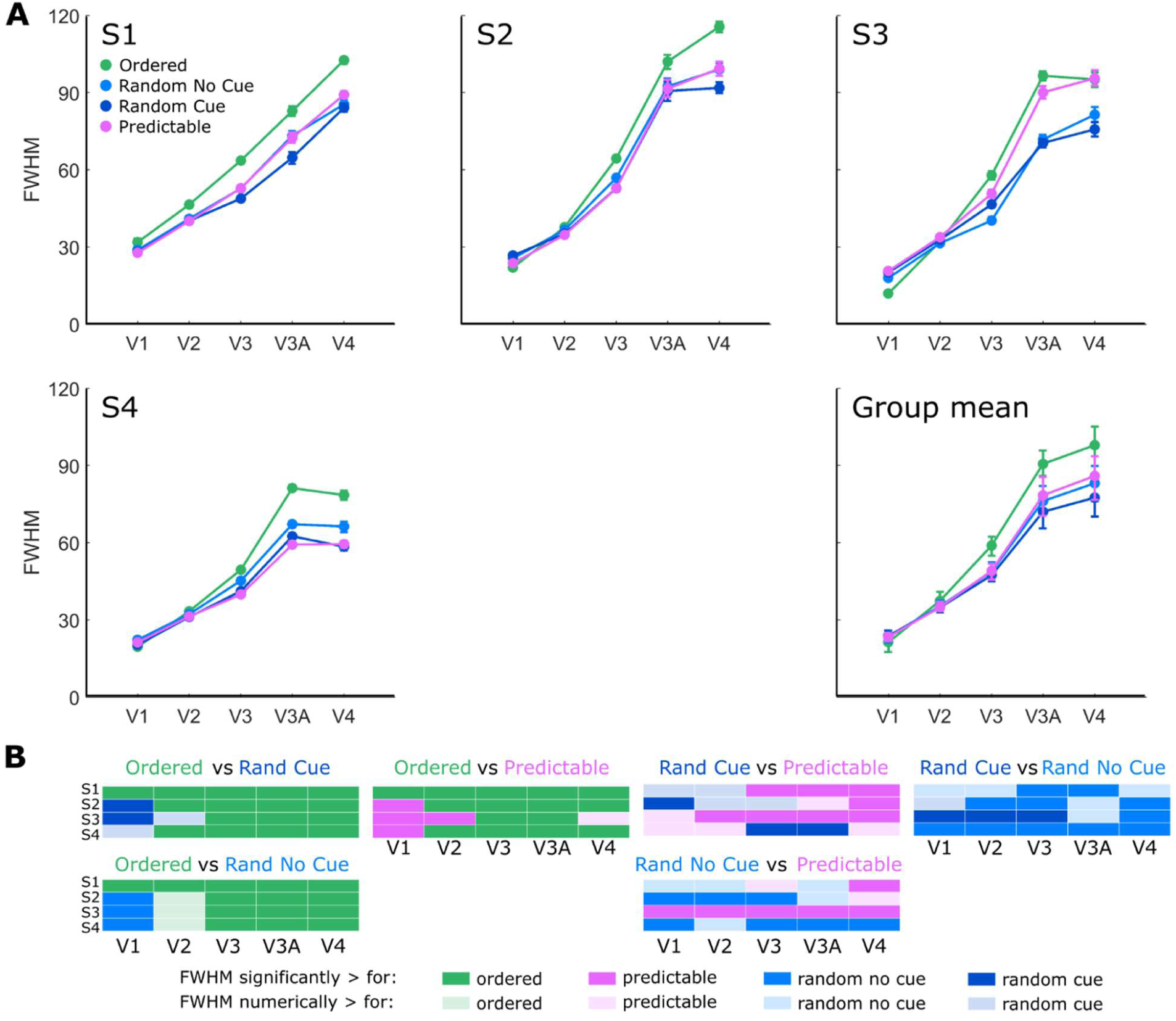
Polar angle tuning width estimates in Experiment 3. A) Individual and group mean FWHM estimates. B) Visualisation of single subject statistics (p<.05 corrected for multiple comparisons).

## Simulations

With the following simulations, we addressed the hypothesis that the discrepancies in tuning width observed for the different mapping sequences in the previous experiments were caused by the spatiotemporal properties of the haemodynamic response (Aquino, Schira, Robinson, Drysdale, & Breakspear, 2012; Kriegeskorte, Cusack, & Bandettini, 2010). To test this possibility, we simulated the BOLD response for Experiments 1 and 3 using the stimBOLD toolbox (https://github.com/KevinAquino/stimBOLD, Aquino, Lacy, Robinson, & Schira, 2015). This toolbox takes a visual input, predicts the cortical neural response that it evokes in early visual cortex (areas V1-V3) and simulates the BOLD response taking into account the poroelastic properties of the brain tissue (Aquino, Robinson, Schira, & Breakspear, 2014; Aquino et al., 2012)

### Materials and methods

#### Stimuli and mapping

We used stimBOLD to simulate the BOLD response in visual areas V1-V3 for the left hemisphere of FreeSurfer average brain (fsaverage) (Benson et al., 2012; Dale et al., 1999; Fischl et al., 1999). We simulated the responses for all the conditions that differed in terms of spatiotemporal structure and stimulus size in the previous experiments. In particular, we selected the ordered, random, and predictable conditions in Experiment 1 – Simulation A – and the ordered and random conditions of Experiment 3 – Simulation B (in both cases we employed the mapping sequences used for participant 4).

The mapping stimulus had the same physical properties adopted in our empirical experiments (max eccentricity = 8.5 dva; wedge size of 45° for Experiment 1 and 6° for Experiment 3). Each location of the visual field was stimulated for 1s, before moving to the next location in the sequence. Two images, selected from the original dataset of natural pictures, alternated every 500 ms.

Six runs for each condition were simulated separately, then Gaussian noise was added to the signal. Gaussian noise was adjusted in order to produce approximately the same signal-to-noise ratio (SNR) across Simulation A and B (Simulation A, SNR_ord_ = .21; SNR_pred_ = .17; SNR_ran_ = .17. Simulation B: SNR_ord_ = .21; SNR_ran_ = .11. We computed the SNR as the ratio between the standard deviation of the signal and the standard deviation of the residuals). The following analyses were performed for the simulated data with and without Gaussian noise.

### Analyses

The signal was z-score normalized and the runs concatenated before modelling the pRF profiles following the same approach used for the empirical data. We focused our analyses on the comparison of FWHM in the different conditions for Simulation A and B considering only those vertices with goodness of fit higher than a critical value based on a fixed p-value (p = 10^−8^, R^2^ > 0.026 for Simulation A and R^2^ > 0.019 for simulation B). We averaged FWHM across eccentricities and ROIs (V1, V2, and V3) separately for each condition and tested their difference with paired t-tests (degrees of freedom corrected for time series correlation).

### Results

The results qualitatively replicated the difference across conditions observed in the empirical data (Figure 11). In Simulation A, the ordered condition resulted in significantly lower estimates of FWHM than the random condition (V1-V3_ord-rand_: t(196.8) = -7.05, p < .001). The predictable condition lead to intermediate results (V1-V3_ord-pred_: t(205.8) = -2.98, p = .003; V1-V3_rand-pred_: t(200.0) = 2.97, p = .003). Crucially, the pattern of results reversed for Simulation B with the ordered condition leading to the significantly higher FWHM estimates than the random one (V1-V3_ord-rand_: t(263.7) = 5.04, p < .001). The analyses of the predicted BOLD response without addition of Gaussian noise produces an inversion of the pattern of results for the ordered and predictable conditions in Simulation A (V1-V3_ord-pred_: t(618.0) = 10.72, p < .001; V1-V3_rand-pred_: t(593.0) = 10.72, p < .001). However, the crucial difference between ordered and random conditions is replicated in both simulations (Simulation A. V1-V3_ord-rand_: t(569.8) = -5.06, p < .001. Simulation B. V1-V3_ord-rand_: t(1150.7) = 5.51, p < .001) (Figure 11).

**Figure 11.**
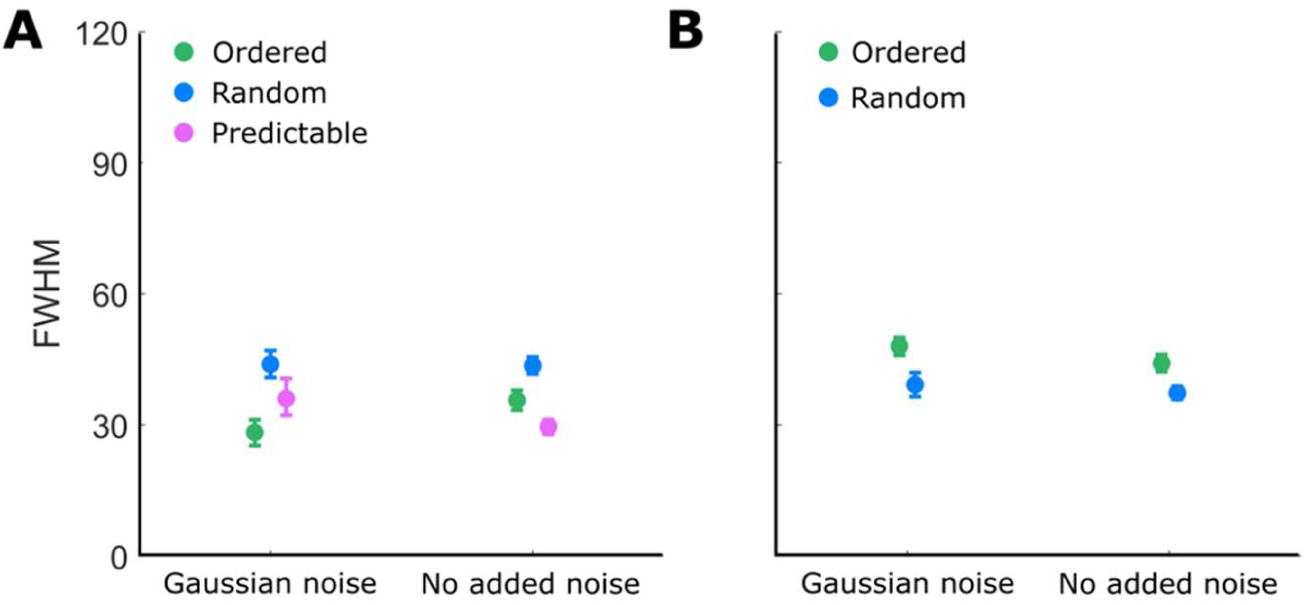
Polar angle tuning width estimates for simulated BOLD responses with and without added Gaussian noise. Error bars in both the individual and group plots represent bootstrapped 95% confidence interval. A) Simulation A replicating the sequence structure and stimulus properties of Experiment 1. B) Simulation B replicating the sequence structure and stimulus properties of Experiment 3.

## Discussion

In this series of experiments, we investigated the reliability and biases of pRF modelling while disambiguating the impact of predictability and spatiotemporal regularities when mapping the visual cortex. We adopted a modified version of the pRF modelling approach (Dumoulin & Wandell, 2008) to estimate the polar angle preference of neural populations in visual cortex and designed mapping sequences characterized by different spatiotemporal structure and different duration.

As reported in previous studies, polar angle estimates were robust across mapping sequences while estimates of pRF size were more volatile (van Dijk et al., 2016). Despite their general robustness, the polar angle estimates in visual areas with larger receptive fields (V3, V3A, V4) were more sensitive to the structure of the mapping sequence when short mapping cycles were adopted. This was particularly evident for the predictable condition in Experiment 1 in which the same short sequence was repeated throughout one run introducing systematic deviations in the measured polar angle estimates.

In all experiments, we observed striking differences in pRF size for ordered and random sequences across the visual areas tested. Interestingly, in Experiment 1 we observed the narrowest tuning widths for ordered mapping sequences, intermediate results for the regular and predictable sequences and the widest tuning width for random sequences. These results are in contrast with previous reports of larger pRF size estimates for ordered sequences (Binda et al., 2013). To test whether such results were a consequence of the anticipation of the attended stimulus, we ran Experiments 2 and 3 where we used a spatial cue to orient attention and matched the spatiotemporal properties of predictable and random sequences. We replicated the findings for ordered and random sequences only and only when a short mapping cycle was employed while the opposite pattern of results emerged for slower designs. Such results argue against an impact of expectations in pRF estimates and suggest that other factors may contribute to these changes in tuning width. One possibility is that fast-paced designs are more susceptible to nonlinear spatiotemporal interactions of responses. For example, centre-surround suppression mechanisms have been suggested to modulate response amplitude when multiple stimuli are presented during mapping, as happens in multi-focal designs (Pihlaja et al., 2008). Alternatively, the rapid stimulation of adjacent regions might induce hemodynamic stealing (Harel, Lee, Nagaoka, Kim, & Kim, 2002), or induce adaptation phenomena (Krekelberg, Boynton, & van Wezel, 2006) hence reduced BOLD signal. Such phenomena could influence pRF size estimates with greater impact on ordered protocols. While these possibilities are intriguing, they cannot easily explain the inversion of the pattern of results observed in our last experiment. Finally, it is possible that active changes in cortical vasculature might be responsible for spatiotemporal nonlinearities in the BOLD response and significantly affect the pRF estimates, particularly with small voxel sizes (Aquino et al., 2012; Kriegeskorte et al., 2010). Our simulations found general support for the last hypothesis indicating a possible mechanism by which the interplay of stimulus properties and mapping sequence can have a measurable impact of pRF estimates.

In our study we did not find any clear evidence that the predictability of stimulus location can significantly bias polar angle or tuning width estimates. This result contradicts previous studies that showed attention can cause both a shift of the preferred location towards the attended location and an increase in pRF size (Kay et al., 2015; Klein et al., 2014; Sheremata & Silver, 2015; van Es et al., 2018; Vo et al., 2017). Such modulation had initially reported only been in the ventral cortex, higher up in the visual cortex (Kay et al., 2015) while more recent evidence suggest that both feature-based and spatial-based spatial attention can induce significant shifts in the response of neurons in areas as early as V1-V3 (van Es et al., 2018). It has been suggested that these changes are functional to increase the precision of the representation of the target at the attended location (Kay et al., 2015). While our design was not tailored to detect systematic changes in polar angle preferences, we hypothesized that the predictability of the mapping sequence would affect the tuning of neuronal responses. The discrepancy between our results and recent observations of attentional effects can be explained by a difference in task requirements among the studies. All former studies manipulated the focus of attention by varying the location at which participants were performing a perceptual task, either at fixation or on the mapping stimulus (Kay et al., 2015; Sheremata & Silver, 2015; van Es et al., 2018). Such demanding tasks required a redistribution of resources at the attended location. On the contrary, in our experiments, our task did not require a fine discrimination and the predictability of stimulus location was not strategically relevant for performing the task. Thus, expectations alone may not dynamically change pRF properties in early visual cortex to a significant extent, as long as there is no computational requirement for that.

Irrespective of the specific sequence, the fitting results described in the current study produced weaker fits than standard mapping approaches (Dumoulin & Wandell, 2008). Several reasons could contribute to these results. First, R^2^ depends considerably on the degrees of freedom. In our experiments, we concatenated the BOLD response in separate runs of the same condition leading to a large number of time points per condition (up to 1818 in Experiment 2) massively increasing the degrees of freedom and generally reducing R^2^ for at statistical significance levels equivalent to other studies. Second, in order to facilitate learning of the predictable sequences, we designed protocols with unusually short cycles in Experiment 1 while long cycles but thin mapping stimuli were employed in the last study. Despite these limitations, we obtained reliable maps in all conditions (Figure 2, Figure 5, Figure 8).

Our study shows that pRF estimates are susceptible to the spatiotemporal properties of the mapping sequence. In particular, ordered and random mapping protocols show different susceptibility to other design choices such as stimulus type and duration of the mapping cycle and can produce significantly different pRF results. Finally, it is worth noting that while ordered sequences are typically preferred for their higher goodness of fit, this is not a guarantee of their robustness. More specifically, the pRF estimates obtained with different sequences, both ordered and random, performed comparably well in predicting the response to different mapping stimuli. To conclude, depending on other design constraints, one should consider which protocol is more suitable for the experimental purposes.

## Acknowledgments

Supported by ERC Starting Grant (310829) to DSS. We thank Kevin Aquino for his support with the use of the BOLD simulation toolbox (stimBOLD).

## Supplementary Materials

### Control analyses and results

In all three experiments, to ensure consistency in fixation stability and task engagement across conditions, we measured target detection accuracy in the dual-task across conditions. Moreover, we measured the mean absolute deviation of eye position from fixation along the x and y axes in degrees of visual angle (dva).

In Experiment 1, we did not observe noticeable differences in participants’ accuracy in performing the dual-task (p(Hit): *M*_*ord*_(*SD*) = .77 (.11), *M*_*pred*_(*SD*) =.74(.14), *M*_*rand*_(*SD*) = .81(.05)) nor in stability of fixation (Absolute deviation from fixation along in dva. Deviation along x axis: *M*_*ord*_(*SD*) = .28(.15); *M*_*pred*_(*SD*) =.24(.11); *M*_*rand*_(*SD*) =.25(.10). Deviation along y axis: *M*_*ord*_(*SD*) = .46(.42), *M*_*pred*_(*SD*) = .41(.33), *M*_*rand*_(*SD*) = .36(.26)) across conditions.

Similarly, in Experiment 2, we did not observe significant differences in participants’ accuracy in the dual-task (p(Hit): *M*_*ord*_(*SD*) = .72 (.16), *M*_*pred*_(*SD*) =.74(.17), *M*_*randNC*_(*SD*) = .68(.08), *M*_*randC*_(*SD*) = .69(.14)) nor in stability of fixation (Absolute deviation from fixation along in dva. Deviation along x axis: *M*_*ord*_(*SD*) = .21(.08), *M*_*pred*_(*SD*) =.19(.06), *M*_*randNC*_(*SD*) =.23(.09), *M*_*randC*_(*SD*) =.21(.09). Deviation along y axis: *M*_*ord*_(*SD*) = .31(.23), *M*_*pred*_(*SD*) = .30(.20), *M*_*randNC*_(*SD*) = .32(.20), *M*_*randC*_(*SD*) =.34(.24) across conditions.

Consistent with the previous experiments, also in Experiment 3, we did not observe significant differences in participants’ accuracy in the dual-task (p(Hit): *M*_*ord*_(*SD*) = .89 (.07), *M*_*pred*_(*SD*) =.86(.15), *M*_*randNC*_(*SD*) = .89(.12), *M*_*randC*_(*SD*) = .93(.06)) nor in stability of fixation (Absolute deviation from fixation along in dva. Deviation along x axis: *M*_*ord*_(*SD*) = .23(.12), *M*_*pred*_(*SD*) =.24(.10), *M*_*randNC*_(*SD*) =.32(.21), *M*_*randC*_(*SD*) =.27(.13). Deviation along y axis: *M*_*ord*_(*SD*) = .31(.22), *M*_*pred*_(*SD*) = .42(.38), *M*_*randNC*_(*SD*) = .54(.61), *M*_*randC*_(*SD*) =.37(.58) across conditions.

**Figure S 1.**
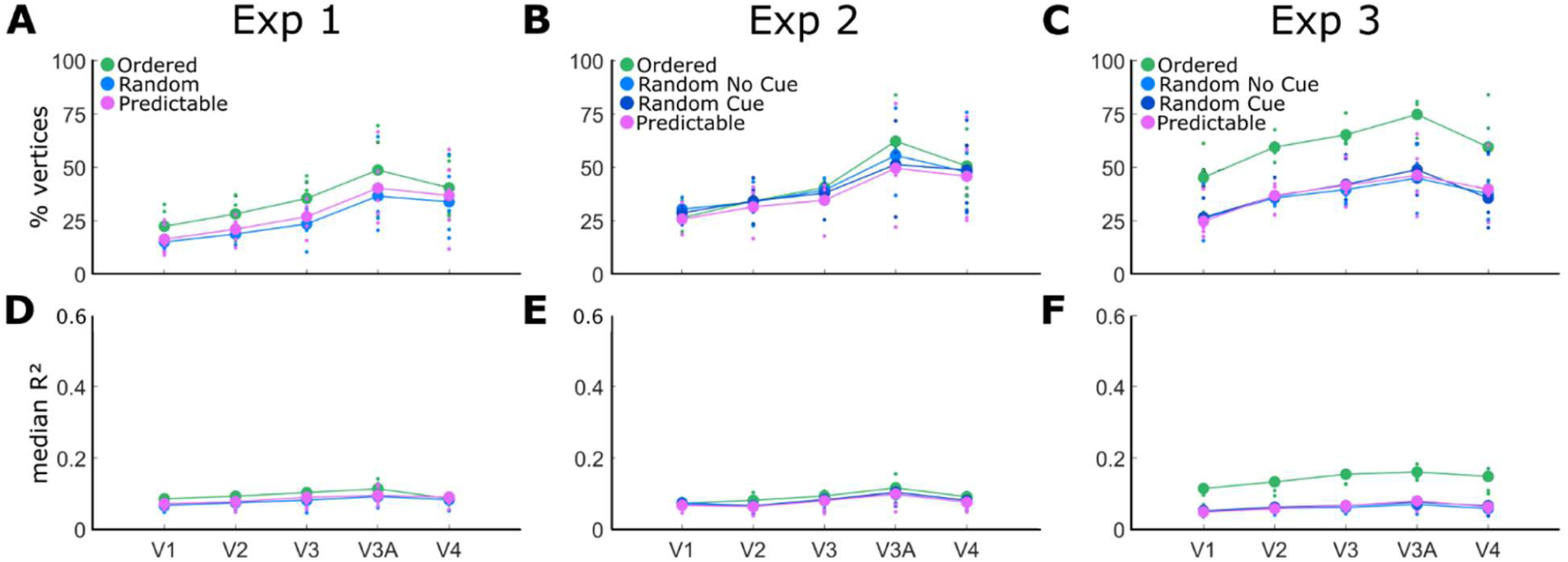
Data quality in Experiments 1-3. (A-C) Percentage of responsive vertices in each visual area and condition tested. (D-F) Median goodness of fit (R^2^). Large dots indicate mean group results and small dots indicate individual participants’ data. Ordered (green), random (blue) and predictable (fuchsia) sequences are displayed for all experiments. In Experiments 2 and 3, the light blue dots and the dark blue dots depict results for the random no-cue and random cue conditions respectively.

**Figure S 2.**
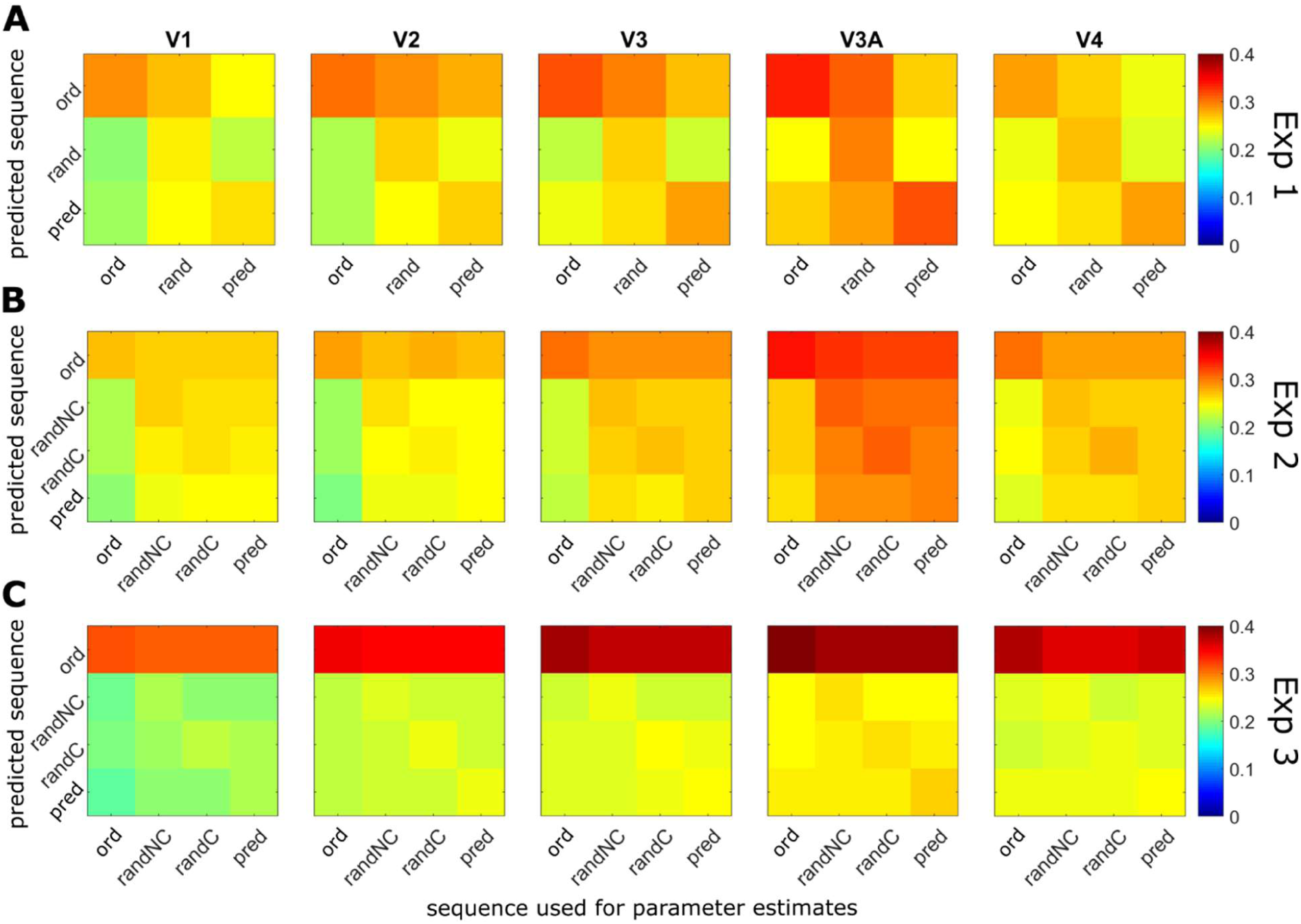
Group mean Pearson correlation matrices of predicted and observed time series in Experiments 1-3. The parameters obtained from fitting each of the mapping sequences (x-axis) are employed to generate time series for each of the conditions in the experiment. The resulting predicted responses are then correlated with the measured time series (y-axis). In all experiments, the ordered sequences are predicted with more accuracy than any of the other sequences irrespective of the parameters used for generating the times series.

## Footnotes

1 One participant performed 15 runs in one single session in Experiment 1.

2 The R^2^ threshold was adjusted to 0.035 for the participant that performed a smaller number of runs in Experiment 1.

3 Here we are not accounting for the degrees of freedom in the pRF model (the p-value would only be marginally different) but the purpose of this procedure is simply to define an objective threshold for the data rather than accurately estimating p-values.

## Bibliography

Alvarez, I., de Haas, B., Clark, C. A., Rees, G., & Schwarzkopf, D. S. (2015). Comparing different stimulus configurations for population receptive field mapping in human fMRI. Frontiers in Human Neuroscience, 9(February), 96. https://doi.org/10.3389/fnhum.2015.00096

Amano, K., Wandell, B. a, & Dumoulin, S. O. (2009). Visual field maps, population receptive field sizes, and visual field coverage in the human MT+ complex. Journal of Neurophysiology, 102(5), 2704–2718. https://doi.org/10.1152/jn.00102.2009

Anderson, E. J., Tibber, M. S., Schwarzkopf, D. S., Shergill, S. S., Fernandez-Egea, E., Rees, G., & Dakin, S. C. (2016). Visual population receptive fields in people with schizophrenia have reduced inhibitory surrounds. The Journal of Neuroscience, 37(6), 3620–15. https://doi.org/10.1523/JNEUROSCI.3620-15.2016

Aquino, K. M., Lacy, T. C., Robinson, P. A., & Schira, M. M. (2015). Using models to design fMRI experiments - not just fit data. In 21st Annual Meeting of the Organization for Human Brain Mapping.

Aquino, K. M., Robinson, P. A., Schira, M. M., & Breakspear, M. (2014). Deconvolution of neural dynamics from fMRI data using a spatiotemporal hemodynamic response function. NeuroImage, 94, 203–215. https://doi.org/10.1016/j.neuroimage.2014.03.001

Aquino, K. M., Schira, M. M., Robinson, P. A., Drysdale, P. M., & Breakspear, M. (2012). Hemodynamic traveling waves in human visual cortex. PLoS Computational Biology, 8(3). https://doi.org/10.1371/journal.pcbi.1002435

Benson, N. C., Butt, O. H., Datta, R., Radoeva, P. D., Brainard, D. H., & Aguirre, G. K. (2012). The retinotopic organization of striate cortex is well predicted by surface topology. Current Biology, 22(21), 2081–2085. https://doi.org/10.1016/j.cub.2012.09.014

Binda, P., Thomas, J. M., Boynton, G. M., & Fine, I. (2013). Minimizing biases in estimating the reorganization of human visual areas with BOLD retinotopic mapping. Journal of Vision, 13(7), 1–16. https://doi.org/10.1167/13.7.13

Brainard, D. H. (1997). The Psychophysics Toolbox. Spatial Vision, 10, 433–436. https://doi.org/10.1163/156856897X00357

Breuer, F. A., Blaimer, M., Heidemann, R. M., Mueller, M. F., Griswold, M. A., & Jakob, P. M. (2005). Controlled aliasing in parallel imaging results in higher acceleration (CAIPIRINHA) for multi-slice imaging. Magnetic Resonance in Medicine, 53(3), 684–691. https://doi.org/10.1002/mrm.20401

Dale, A. M., Fischl, B., & Sereno, M. I. (1999). Cortical surface-based analysis: I. Segmentation and surface reconstruction. NeuroImage, 9(2), 179–194. https://doi.org/10.1006/nimg.1998.0395

de Haas, B., Schwarzkopf, D. S., Anderson, E. J., & Rees, G. (2014). Perceptual load affects spatial tuning of neuronal populations in human early visual cortex. Current Biology, 24(2), R66–R67. https://doi.org/10.1016/j.cub.2013.11.061

Dekker, T. M., Schwarzkopf, D. S., de Haas, B., Nardini, M., & Sereno, M. I. (2017). Population receptive field tuning properties of visual cortex during childhood. BioRxiv:2132108. https://doi.org/10.1101/213108

Dekker, T. M., Schwarzkopf, D. S., de Haas, B., Nardini, M., & Sereno, M. I. (2019). Population receptive field tuning properties of visual cortex during childhood. Developmental Cognitive Neuroscience, (July 2018), 100614. https://doi.org/10.1016/j.dcn.2019.01.001

Dumoulin, S. O., & Knapen, T. (2018). How Visual Cortical Organization Is Altered by Ophthalmologic and Neurologic Disorders. Annual Review of Vision Science, 4(1), annurev-vision-091517-033948. https://doi.org/10.1146/annurev-vision-091517-033948

Dumoulin, S. O., & Wandell, B. A. (2008). Population receptive field estimates in human visual cortex. NeuroImage, 39(2), 647–660. https://doi.org/10.1016/j.neuroimage.2007.09.034

Ekman, M., Kok, P., & de Lange, F. P. (2017). Time-compressed preplay of anticipated events in human primary visual cortex. Nature Communications, 8(May), 15276. https://doi.org/10.1038/ncomms15276

Engel, S. A., Rumelhart, D. E., Wandell, B. A., Lee, A. T., Glover, G. H., Chichilnisky, E.-J., & Shadlen, M. N. (1994). fMRI of human visual cortex. Nature. https://doi.org/10.1038/369525a0

Fischl, B., Sereno, M. I., & Dale, A. M. (1999). Cortical surface-based analysis: II. Inflation, flattening, and a surface-based coordinate system. NeuroImage, 9(2), 195–207. https://doi.org/10.1006/nimg.1998.0396

Gomez, J., Natu, V., Jeska, B., Barnett, M., & Grill-Spector, K. (2018). Development differentially sculpts receptive fields across early and high-level human visual cortex. Nature Communications, 9(1), 788. https://doi.org/10.1038/s41467-018-03166-3

Harel, N., Lee, S. P., Nagaoka, T., Kim, D. S., & Kim, S. G. (2002). Origin of negative blood oxygenation level-dependent fMRI signals. Journal of Cerebral Blood Flow and Metabolism, 22(8), 908–917. https://doi.org/10.1097/00004647-200208000-00002

Harvey, B. M., & Dumoulin, S. O. (2011). The relationship between cortical magnification factor and population receptive field size in human visual cortex: constancies in cortical architecture. J Neurosci, 31(38), 13604–13612. https://doi.org/10.1523/JNEUROSCI.2572-11.2011

Kastner, S., Pinsk, M. A., De Weerd, P., Desimone, R., & Ungerleider, L. G. (1999). Increased activity in human visual cortex during directed attention in the absence of visual stimulation. Neuron, 22(4), 751–61. https://doi.org/http://dx.doi.org/10.1016/S0896-6273(00)80734-5

Kay, K. N., Weiner, K. S., & Grill-Spector, K. (2015). Attention reduces spatial uncertainty in human ventral temporal cortex. Current Biology, 25(5), 595–600. https://doi.org/10.1016/j.cub.2014.12.050

Klein, B. P., Harvey, B. M., & Dumoulin, S. O. (2014). Attraction of position preference by spatial attention throughout human visual cortex. Neuron, 84(1), 227–237. https://doi.org/10.1016/j.neuron.2014.08.047

Krekelberg, B., Boynton, G. M., & van Wezel, R. J. A. (2006). Adaptation: from single cells to BOLD signals. Trends in Neurosciences, 29(5), 250–256. https://doi.org/10.1016/j.tins.2006.02.008

Kriegeskorte, N., Cusack, R., & Bandettini, P. (2010). NeuroImage How does an fMRI voxel sample the neuronal activity pattern : Compact-kernel or complex spatiotemporal filter ? NeuroImage, 49(3), 1965–1976. https://doi.org/10.1016/j.neuroimage.2009.09.059

Moutsiana, C., de Haas, B., Papageorgiou, A., van Dijk, J. A., Balraj, A., Greenwood, J. A., & Schwarzkopf, D. S. (2016). Cortical idiosyncrasies predict the perception of object size. Nature Communications, 7, 1–25. https://doi.org/10.1101/026989

Pelli, D. G. (1997). The VideoToolbox software for visual psychophysics: transforming numbers into movies. Spatial Vision, 10, 437–442. https://doi.org/10.1163/156856897X00366

Pihlaja, M., Henriksson, L., James, A. C., & Vanni, S. (2008). Quantitative multifocal fMRI shows active suppression in human V1. Human Brain Mapping, 29(9), 1001–1014. https://doi.org/10.1002/hbm.20442

Schwarzkopf, D. S., Anderson, E. J., de Haas, B., White, S. J., & Rees, G. (2014). Larger Extrastriate Population Receptive Fields in Autism Spectrum Disorders. Journal of Neuroscience, 34(7), 2713–2724. https://doi.org/10.1523/JNEUROSCI.4416-13.2014

Sereno, M. I., Dale, a M., Reppas, J. B., Kwong, K. K., Belliveau, J. W., Brady, T. J., … Tootell, R. B. H. (1995). Borders of Multiple Visual Areas in Humans Revealed by Functional Magnetic Resonance Imaging Borders of Multiple Visual Areas in Humans Revealed by Functional Magnetic Resonance Imaging. Science, 268(5212), 889–893.

Sereno, M. I., McDonald, C. T., & Allman, J. M. (1994). Analysis of Retinotopic Maps in Extrastriate Cortex. Cerebral Cortex, 4(6), 601–620. https://doi.org/10.1093/cercor/4.6.601

Sheremata, S. L., & Silver, M. A. (2015). Hemisphere-Dependent Attentional Modulation of Human Parietal Visual Field Representations. Journal of Neuroscience, 35(2), 508–517. https://doi.org/10.1523/JNEUROSCI.2378-14.2015

Silson, E. H., Reynolds, R. C., Kravitz, D. J., & Baker, C. I. (2018). Differential sampling of visual space in ventral and dorsal early visual cortex. The Journal of Neuroscience, 38(9), 2717–17. https://doi.org/10.1523/JNEUROSCI.2717-17.2018

Silva, M. F., Brascamp, J. W., Ferreira, S., Castelo-Branco, M., Dumoulin, S. O., & Harvey, B. M. (2018). Radial asymmetries in population receptive field size and cortical magnification factor in early visual cortex. NeuroImage, 167(September 2016), 41–52. https://doi.org/10.1016/j.neuroimage.2017.11.021

Smittenaar, C. R., Macsweeney, M., Sereno, M. I., & Schwarzkopf, D. S. (2016). Does Congenital Deafness Affect the Structural and Functional Architecture of Primary Visual Cortex? The Open Neuroimaging Journal, 10(16), 1–19. https://doi.org/10.2174/1874440001610010001

Thomas, J. M., Huber, E., Stecker, G. C., Boynton, G. M., Saenz, M., & Fine, I. (2015). Population receptive field estimates of human auditory cortex. NeuroImage, 105, 428–439. https://doi.org/10.1016/j.neuroimage.2014.10.060

van Dijk, J. A., de Haas, B., Moutsiana, C., & Schwarzkopf, D. S. (2016). Intersession reliability of population receptive field estimates. NeuroImage, 143, 293–303. https://doi.org/10.1016/j.neuroimage.2016.09.013

van Es, Theeuwes, J., & Knapen, T. (2018). Spatial sampling in human visual cortex is modulated by both spatial and feature-based attention.

Vanni, S., Henriksson, L., & James, A. C. (2005). Multifocal fMRI mapping of visual cortical areas. NeuroImage, 27(1), 95–105. https://doi.org/10.1016/j.neuroimage.2005.01.046

Vo, V. A., Sprague, T. C., & Serences, J. T. (2017). Spatial Tuning Shifts Increase the Discriminability and Fidelity of Population Codes in Visual Cortex. The Journal of Neuroscience, 37(12), 3386–3401. https://doi.org/10.1523/JNEUROSCI.3484-16.2017

Wandell, B. A., & Winawer, J. (2015). Computational neuroimaging and population receptive fields. Trends in Cognitive Sciences, 19(6), 349–357. https://doi.org/10.1016/j.tics.2015.03.009

Yildirim, F., Carvalho, J., & Cornelissen, F. W. (2018). A second-order orientation-contrast stimulus for population-receptive-field-based retinotopic mapping. NeuroImage, 164(May), 183–193. https://doi.org/10.1016/j.neuroimage.2017.06.073

Zeidman, P., Silson, E. H., Schwarzkopf, D. S., Baker, C. I., & Penny, W. (2018). Bayesian population receptive field modelling. NeuroImage, 180(September), 173–187. https://doi.org/10.1016/j.neuroimage.2017.09.008

Zuiderbaan, W., Harvey, B. M., & Dumoulin, S. O. (2012). Modeling center – surround configurations in population receptive fields using fMRI. Journal of Vision, 12(3), 1–15. https://doi.org/10.1167/12.3.10.Introduction

